# Cause and consequences of genome duplication in haploid yeast populations

**DOI:** 10.1101/247320

**Authors:** Kaitlin J. Fisher, Sean W. Buskirk, Ryan C. Vignogna, Daniel A. Marad, Gregory I. Lang

## Abstract

Whole genome duplications (WGD) represent important evolutionary events that shape future adaptation. WGDs are known to have occurred in the lineages leading to plants, fungi, and vertebrates. Changes to ploidy level impact the rate and spectrum of beneficial mutations and thus the rate of adaptation. Laboratory evolution experiments initiated with haploid *Saccharomyces cerevisiae* cultures repeatedly experience WGD. We report recurrent genome duplication in 46 haploid yeast populations evolved for 4,000 generations. We find that WGD confers a fitness advantage, and this immediate fitness gain is accompanied by a shift in genomic and phenotypic evolution. The presence of ploidy-enriched targets of selection and structural variants reveals that autodiploids utilize adaptive paths inaccessible to haploids. We find that autodiploids accumulate recessive deleterious mutations, indicating an increased capacity for neutral evolution. Finally, we report that WGD results in a reduced adaptation rate, indicating a trade-off between immediate fitness gains and long term adaptability.

## INTRODUCTION

The natural life cycle of budding yeast alternates between haploid and diploid phases. Both ploidies can be stably propagated asexually through mitotic division. Both theory and experimental work show that haploids adapt faster than diploids, likely due to recessive beneficial mutations (Orr and Otto 1994; Zeyl, Vanderford, Carter 2003). Curiously, however, repeated attempts at evolving experimental haploid populations have resulted in recurrent whole genome duplications yielding populations of autodiploids (Gerstein *et al.* 2006; Hong and Gresham 2014; Voordeckers *et al.* 2015); see Table 1 for additional). Proposed explanations of this phenomenon include artifacts of strain construction (Venkataram *et al.* 2016), unintended mating events (Voordeckers *et al.* 2015), and an adaptive advantage of diploidy (Gerstein *et al.* 2006).

**Table 1:** Observations of autodiploidy in experimental studies.

| Study | Propagation | Evolution medium | Strain background | Mating-type |
| --- | --- | --- | --- | --- |
| Current study | Batch culture, unshaken | YPD | W303 | *MATa* & *MATα* |
| Kosheleva and Desai 2017 | Batch culture, unshaken | YPD | Sk1-W303 hybrid | *MATa* & *MATα* |
| Gorter 2017 | Batch culture, shaken | YPD with heavy metals | BY4743 | *MATa* |
| Venkataram *et al.* 2016 | Batch culture, shaken | Carbon limited glucose | BY4709 | *MATa* |
| Voordeckers *et al.* 2015 | Turbidostat | 6-12% EtOH glucose | S288c derivative | *MATα* |
| Hong and Gresham 2014 | Chemostat | Nitrogen limited glucose | S288c derivative | *MATa* |
| Oud *et al.* 2013 | Anerobic batch culture in sequential bioreactor | 1:1 glucose/galactose | CEN.PK113-7D | *MATa* |
| Gerstein *et al.* 2006 | Batch culture, shaken | YPD | SM2185 | *MATa* |

Whole genome duplication (WGD) in asexual haploid populations could provide a fitness advantage in several different ways. Cell size scales with DNA content in yeast (Gregory 2001), and increased cell size may facilitate more rapid metabolism and increased growth rate. Indeed, increased cell volume has been reported in laboratory-evolved microbial populations (Lenski and Travisano 1994). Gene expression patterns also vary with ploidy (Galitski *et al.* 1999), and diploid-specific gene regulation may be optimal. “Ploidy drive” has been used to describe the phenomenon by which ploidy changes in evolving fungi favor restoration of the historical ploidy state (Gerstein *et al.* 2017). Natural *Saccharomyces cerevisiae* isolates are typically diploid (Liti 2015) and occasionally polyploid (Ezov *et al.* 2006). If most selection has occurred on these higher ploidy states, then gene regulation and cell physiology of diploids should be better optimized relative to haploids.

Despite the recurrence of diploidization events in haploid-founded yeast lineages, the nature of the fitness advantage of diploidy remains unclear. Some studies detect a fitness benefit (Gorter *et al.* 2017; Venkataram *et al.* 2016), while no advantage is detected in others (Gerstein and Otto 2011; Hong and Gresham 2014). A survey of the effect of ploidy on growth rate in otherwise isogenic strains indicates that the benefit of ploidy varies across conditions and optimal ploidy states are contingent on environment (Zörgö *et al.* 2013). In environments where duplication does not confer a direct fitness advantage, it may afford indirect benefits that are then themselves acted upon by selection. Diploidy may protect evolving lineages from purifying selection through buffering the effects of deleterious recessive mutations. Indeed, 15% of viable single gene deletions in haploids exhibit growth defects in rich media, while 97% of heterozygous gene deletions show no detectable phenotype in the absence of perturbation (Deutschbauer *et al.* 2005). This “masking” hypothesis also has experimental support from mutagenesis studies (Mable and Otto 2001), and this effect could be advantageous in populations in which the deleterious mutation rate is sufficiently high.

Autodiploids could invade haploid populations due to increased access to beneficial mutations. Ploidy-dependent mutations are known to arise in experimental evolution (Gerstein 2013; Marad and Lang 2017), and a favorable shift in the distribution of fitness effects may follow genome duplication. Structural variants - deletions, amplifications, and translocations - have repeatedly been shown to be adaptive in experimentally evolving yeast populations (Dunham *et al.* 2002; Gresham *et al.* 2008). Diploids have a greater tendency to form copy number variants (CNVs), especially large deletions (Zhang *et al.* 2013). Likewise, aneuploidies accumulate at a significantly higher rate in diploids in the absence of selection (N. Sharp, *personal communication,* Aug. 2017). If structural variants are more frequent, more variable, and more tolerable in diploids, genome duplication may enable access to novel adaptive paths. Given the repeated observation of displacement of haploids by diploids (Table 1), and the absence of clear evidence for instantaneous fitness advantages of isogenic diploidy that is broadly applicable across experiments, it is possible that selection for and maintenance of diploidy is a complex process involving both direct selection on ploidy state and second order selection, or selection for indirect fitness benefits associated with higher ploidy.

Here we show recurrent WGD in 46 haploid-founded populations during 4,000 generations of laboratory evolution in rich media. We track the dynamics of genome duplication across the haploid-founded populations, revealing that autodiploids fix by generation 1,000 in all 46 populations. Competitive fitness assays show that WGD provides a 3.6% fitness benefit in the selective environment. We find that the immediate fitness gain is accompanied by a loss of access to recessive beneficial mutations. As a consequence, the rate of adaptation of autodiploids slows. Sequencing of the evolved genomes indicates that autodiploids have increased access to structural variants and largely utilize a different spectrum of mutations to adapt compared to haploids. Finally, we show that autodiploids are buffered from the effects of recessive deleterious mutations, consistent with a long-term benefit to maintaining a diploid genome and loss of redundancy following WGD.

## RESULTS

### Sequenced genomes indicate early and recurrent fixation of autodiploids

Two clones were sequenced from each of 46 haploid-founded populations after 4,000 generations of evolution, revealing over 5,100 *de novo* mutations distributed uniformly across the genome, representing the largest dataset of mutations identified in *S. cerevisiae* experimental evolution to date (**Fig. S1**; **Dataset 1**). Mutations are normally distributed across clones (one-sample Kolmogorov-Smirnov test, α=0.05) with a mean of 91 ± 20 (**Fig. S2A**). Most mutations in the sequenced clones were called at ~0.5 (implying heterozygosity), a surprising result given that the populations were founded by a haploid ancestor. Recurrent WGD events were suspected given that each clone maintained its ancestral mating type allele. Further, this hypothesis of WGD was supported by the observation that clones are not heterozygous at polymorphic sites that differed between the *MAT***a** and *MAT*α ancestors. Finally, evolved autodiploids are mating competent, pointing to duplication of haploid genotypes.

### Autodiploids are detected early, sweep quickly, and exhibit a fitness advantage

We determined the fitness effect of genome duplication by directly competing *MAT***a**/**a** autodiploids against an otherwise isogenic haploid *MAT***a** reference. To control for possible artifacts of construction, we independently constructed and competed 10 *MAT***a**/**a** diploids. All 10 *MAT***a**/**a** autodiploid reconstructions exhibit a relative fitness advantage significantly higher than a control haploid strain (Welch’s t-test, *t*=16.28 *df* =19*, p*<.001). Genome duplication alone in the absence of any other variation provides a mean fitness benefit of 3.6% in these experimental conditions (Fig. 1A).

**Fig. 1.**
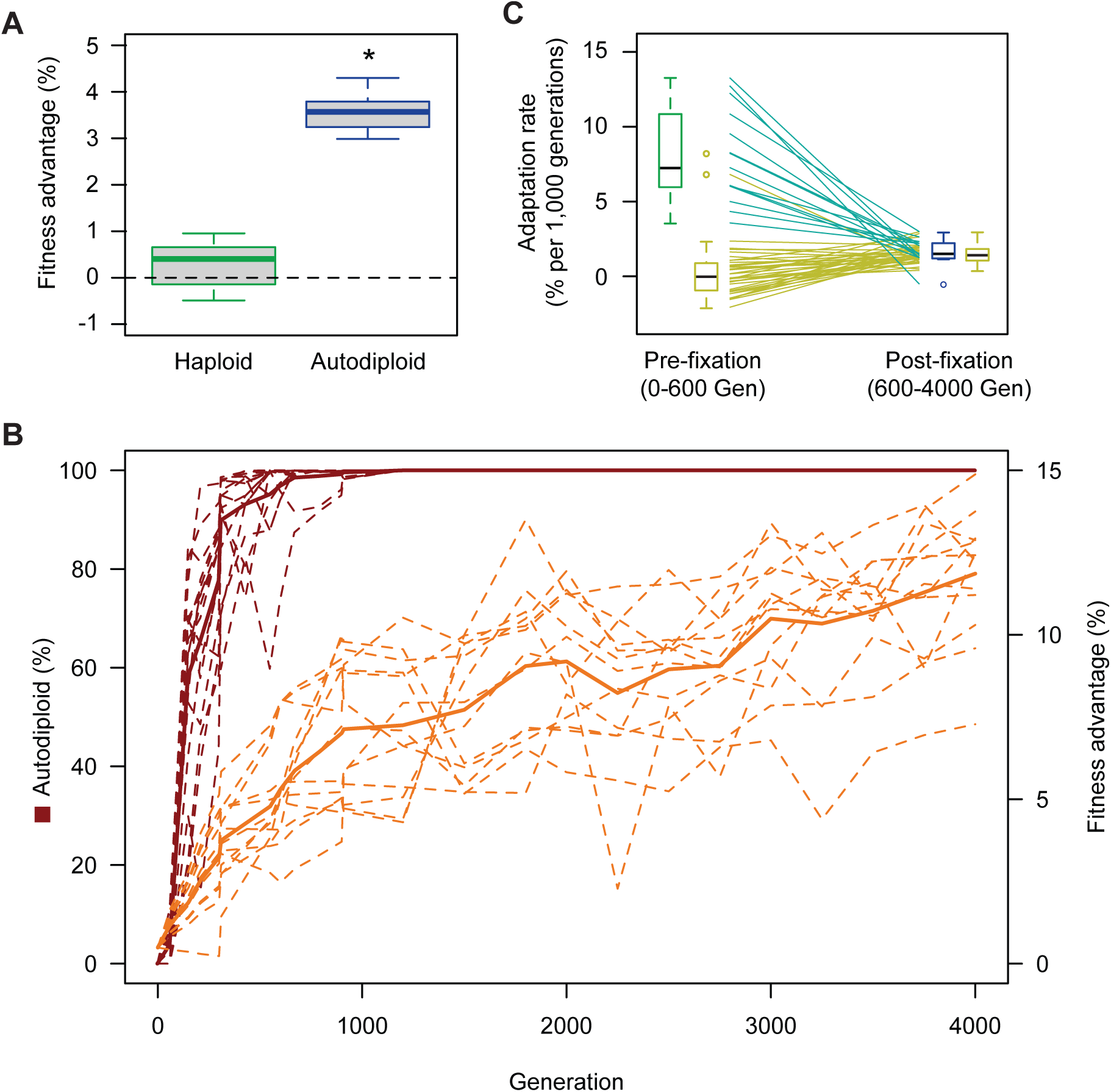
Autodiploids sweep through haploid populations due to a direct fitness advantage. A) *MAT****a****/****a*** diploids have a mean relative fitness advantage of 3.6% when competed against a haploid reference strain. Ten *MAT****a****/****a*** diploids clones were constructed independently. Box plots reflect mean fitness of each clone. Autodiploids and control haploids were competed against the same haploid reference. Asterisk (*) indicates *p*<0.001 (Welch’s t-test, *df*=18.268) B)Autodiploid frequency (red) and fitness advantage (orange) for focal populations (dashed lines). Solid lines indicate mean autodiploid frequency for 16 populations and mean fitness advantage for 13 populations. C) Haploid-founded populations demonstrate significantly higher rates of adaptation until autodiploids fix in the haploid-founded populations. From that point forward, haploid-founded (autodiploids) and diploid-founded populations adapt at the same rate. Lines indicate paired data points from the same population (teal: haploid-founded, yellow: diploid-founded). For each haploid-founded population, adaptation rate was calculated before and after autodiploid fixation, which occured on average at generation 600. Adaptation rates for diploid-founded populations were calculated from Gen 0-600 and Gen 600-4000.

To determine the timing of duplication events, we performed time-course DNA content staining on cryoarchived samples for 16 populations (8 of each mating-type). Autodiploids arise quickly in all 16 populations, fixing by generation 1,000 in all but 2 populations (Fig. 1B, **Fig. S3**, **Fig. S4**). Diploids are present at 2% − 11% in 11/16 populations at generation 60, the earliest time point available for assay. Some populations appear to show clonal interference by fit haploids, with autodiploid fractions briefly decreasing between some time points. Aside from such slight variations, patterns of emergence and spread of autodiploids display show similar dynamics for all 16 populations examined.

We examined whether the degree of parallelism observed in ploidy dynamics can be attributed to ancestral ploidy polymorphisms present at the onset of the experiment. Three lines of evidence support the independent origin of autodiploidy in this experiment. First, the cultures were initiated from two starting strains (*MAT***a** and *MAT*α). There is no significant difference in autodiploid frequency between mating-types at any generation (**Fig. S3**), meaning if autodiploids did, in fact, arise in both independent inoculating cultures, they would have had to achieve roughly the same frequency, which is highly unlikely. Second, no diploids were detected by DNA content staining in any populations at generation 0, indicating autodiploids were not present in the inocula above our detection limit of 1%. Third, computational simulations show that low frequency autodiploids are insufficient to explain the recurrent observation of autodiploid fixation events in all 46 replicate populations. Autodiploids with a 3.6% fitness advantage starting at a frequency of 0.01, the highest frequency we modeled, have a probability of fixation an a given population of 0.88 and therefore the chance of fixation in all 46 populations would be 2.5 × 10^−3^ (**Fig. S5**). Taken together, this argues that, while ancestral autodiploids may have swept in some populations, ancestral ploidy variation is insufficient to explain autodiploid fixation in all 46 populations. Therefore independent, parallel WGD events during the evolution experiment are necessary to explain the recurrent fixation reported here.

### Autodiploids adapt more slowly than haploids

To examine how the shift to diploidy impacted the dynamics of adaptive evolution, we measured population fitness for all populations at ~300-generation intervals. Mean time-course fitness estimates show a change in slope following 1,000 generations. This corresponds roughly to the time that autodiploids have fixed in most focal populations and are high frequency in the remaining populations (Fig. 1B). We compared the rate of adaptation before and after the fixation of diploids in 13 focal populations for which quality fitness data was available. Because many factors, including epistasis, could explain a change in adaptation rate over time, we used a repeated measures ANOVA to compare the effect of ploidy on adaptation rate using time-course fitness data from diploid-founded populations that were evolved in parallel (Marad and Lang 2017) (Fig. 1C). The interaction of founding ploidy and generation has a significant effect (*F*(1, 49)=78.04, *p*<.001, *η*_*p*_^*2*^ = 0.614). Post hoc comparisons using a Bonferroni correction indicate that rates of adaptation are significantly higher in haploid-founded populations than diploids (*p*<.001), and that adaptation rate does not differ once autodiploids fix (*p*=.38). These data corroborate previous findings regarding the effect of ploidy on adaption rate (Gerstein *et al.* 2011; Marad and Lang 2017) and show that autodiploidy provides an immediate fitness gain at the expense of slowing subsequent adaptation.

### Autodiploid genomes harbor autodiploid specific mutations

Duplication of a haploid genome affects both cell physiology and the phenotypic consequences of new mutations. Therefore, the selective pressure on a gene may vary depending on ploidy state. To understand how genome evolution is driving adaptation in the autodiploid populations, we utilize a recurrence approach that accounts for both the number of mutations observed in a gene and the expectation that the observed number of mutations of a given gene occurred by chance alone controlling for gene length. The resulting probabilities were used to identify 20 common genic targets of selection (Fig. 2A). There is a median of 4 recurrent targets per clone with only 1 population containing no common target mutations. GO-component term analysis indicates common targets are enriched for genes whose protein products localize to the cell periphery (*p* = 0.001). Cell periphery targets include *CCW12* and *KRE6*, which both appear to be under extremely strong selective pressure when using the probability metric as a proxy for strength of selection. Interestingly, a tRNA gene, tL(GAG)G, was also identified as a common target of selection (**Fig. S6**). This is the first evidence of adaptive tRNA mutations in laboratory yeast evolution.

**Fig. 2.**
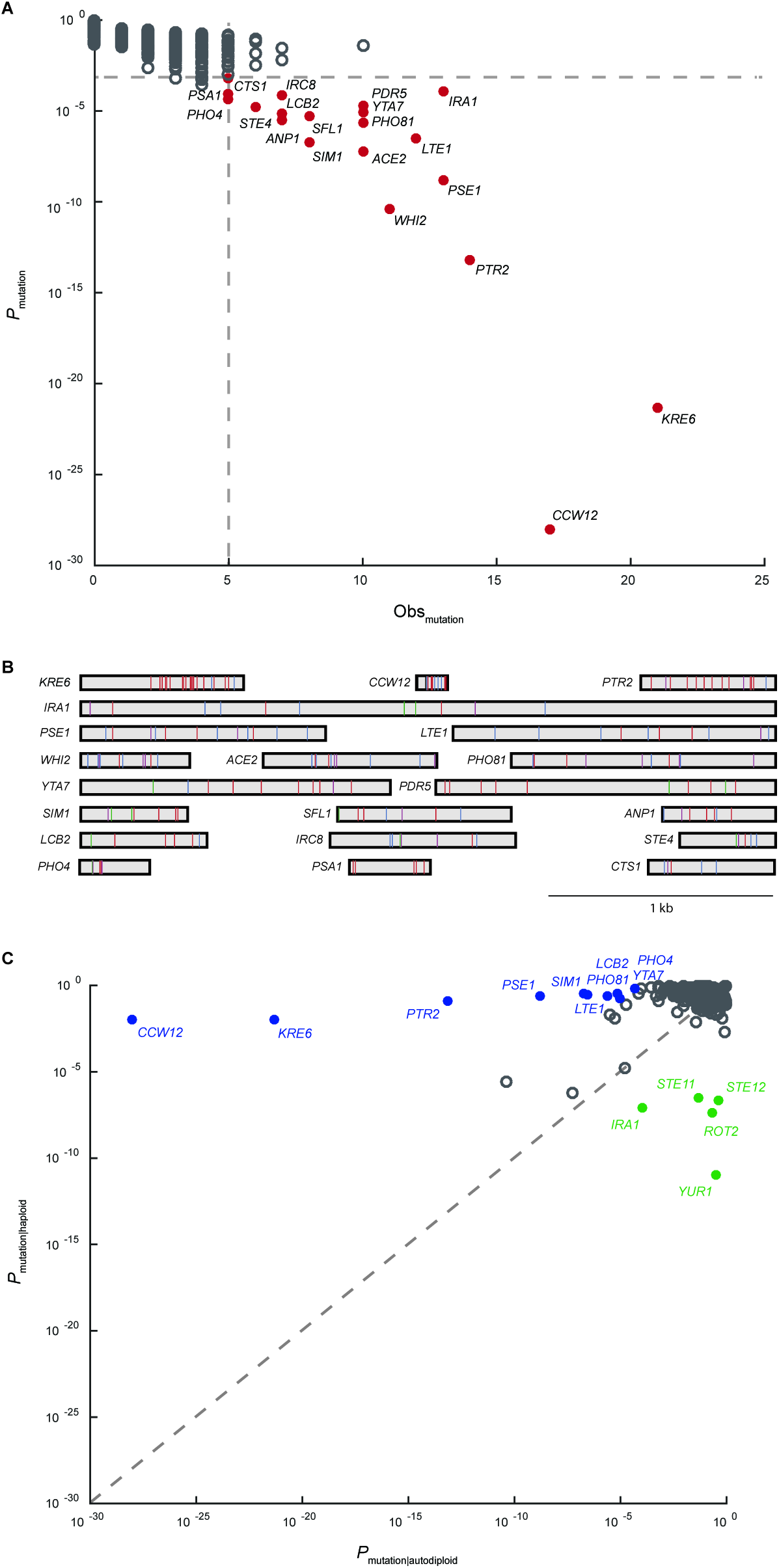
Common targets of selection and ploidy-enriched genes. A) Plotted on the x-axis is the observed number of coding sequence (CDS) mutations in each of the 5800 genes in the S288c reference genome. On the y-axis is the probabilty that the observed number of CDS mutations in each gene occurred by chance. Common targets of selection (solid red circles) are genes with 5 or more CDS mutations and corresponding probability of less than 0.1%. B) Shown are all 188 mutations across the 20 common targets of selection. Genes are represented as rectangles and labelled by gene name. Mutations are colored by type: frameshift-purple, nonsense-blue, missense-red, synonymous-green, other-black. Both homozygous and heterozygous mutations are shown. C) Plotted is the probability that the observed number of CDS mutations in a gene occurred by chance in haploid populations versus autodiploid populations. Genes were considered ploidy-enriched if the ratio of probabilities was greater than 10^5^. Haploid-enriched genes are indicated by solid green circles and autodiploid-enriched genes as solid blue circles.

To better understand the functional basis of adaptation, we examined the distribution of mutations within each gene (Fig. 2B). Three broad patterns emerge. First, we observe selection for loss-of-function alleles, e.g. 9 of 11 mutations in *WHI2* are high impact (frameshift or nonsense). Adaptive loss-of-function alleles are common in experimental microbial evolution (Cooper *et al.* 2001; Kvitek and Sherlock 2013; Venkataram *et al.* 2016). We also observe selection for change-of-function alleles. For example, only missense and synonymous mutations are seen in *PDR5*. Finally, we observe mutations in common targets that cluster within specific domains. This is illustrated by the clustering of mutations in the C-terminus of both *KRE6* (n=21) and *STE4* (n=6).

We compared the common targets of selection identified in autodiploid clones to those identified with the same approach in a comparable haploid dataset (Lang *et al.* 2013) (**Fig. S7**). We identify several haploid‐ and autodiploid-enriched targets (Fig. 2C). Ploidy-enriched targets include genes mutated more often in one ploidy (e.g. *CCW12* and *KRE6* in autodiploids; *YUR1 and ROT2* in haploids) or exclusively in one ploidy (e.g. *PHO81*, *YTA7*, *IRC8* in autodiploids; *STE12* in haploids).

### Loss of heterozygosity hotspots occur on Chromosomes VII and XV

Homozygous mutations, while the minority, are common. Clones contain between 0 and 17 homozygous mutations, with an average of 5.4. Homozygous mutations could either represent mutations that arose before duplication events or loss of heterozygosity (LOH) of heterozygous mutations. We find that the homozygous mutations are not distributed randomly throughout the genome; instead, they tend to cluster in particular regions of the genome (Fig. 3). These clusters, located on the right arms of Chr. XII and Chr. XV, account for 55% of all homozygous mutations. This clustering implies that most homozygous variants result from recombination events. By removing homozygous mutations occurring in these regions from analysis, the average number of homozygous mutations per clone drops to 2.4. This indicates that only a few mutations arose in a haploid background. Most genome evolution, therefore occurred after WGD, and thus genome duplications occurred early in the 4,000 generation evolution experiment.

**Fig. 3.**
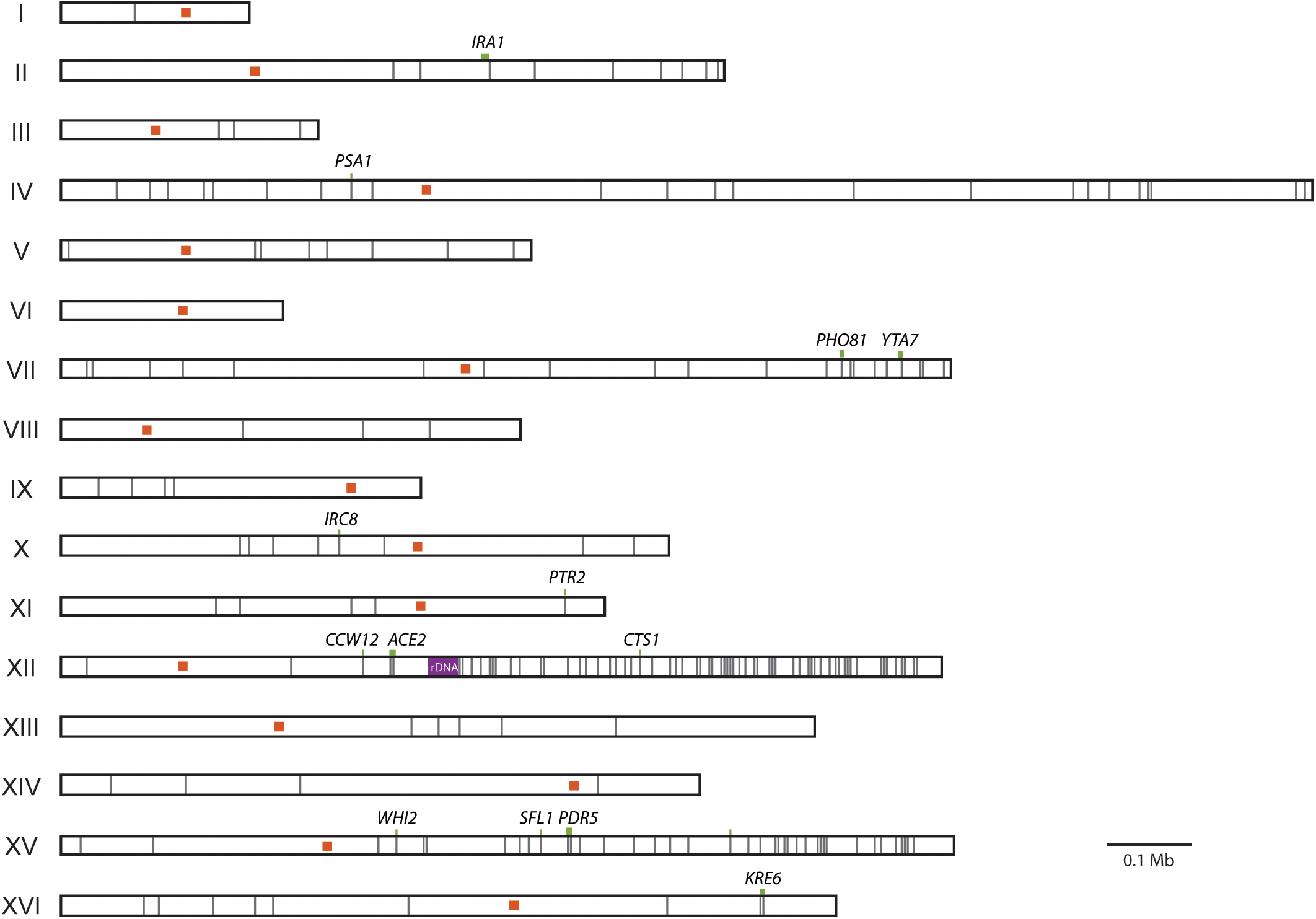
Enrichment of homozygous mutations on the right arms of Chr. XII and Chr. XV. Shown in gray lines are the 256 homozygous mutations detected across the 92 evolved clones. Chromosomes are labeled by Roman numeral. Centromeres are shown as orange squares. Homozygous mutations in common targets of selection are marked by a green line (representing gene length) and labeled by gene name. The ribosomal DNA repeat region of Chr. XII, a known recombination hotspot, is shown in purple.

Mutations in the common targets of selection are observed in various states zygosity. Most genes (12/20) are found mutated in both heterozygous and homozygous states across clones, indicating partial or full dominance of fitness effects. Seven genes only ever contain heterozygous mutations (*ANP1, LCB2, LTE1, PHO4, SIM1, STE4, YTA7*). These mutations are candidates for overdominant effects (Sellis *et al.* 2011). Finally, only one gene, *CTS1*, is never found mutated in a heterozygous state. A reasonable hypothesis would be that the *cts1* mutations are recessive; however, we have previously identified *cts1* mutations in evolved diploid populations and found it to be close to fully dominant (Marad and Lang 2017). Instead, the position of *CTS1* on the right arm of Chr. XII, a LOH hotspot, could explain why it is only observed in a homozygous state (Fig. 3).

### Structural variants are common to autodiploids

In addition to changing the genetic targets of selection, genome duplication permits access to structural variants not accessible to haploid genomes. We analyzed aneuploidies and CNVs in autodiploid genomes as well as previously sequenced haploid populations (Lang *et al.* 2013) (**Figs. 4 & S8; Datasets 2 & 3**). Two types of aneuploidies are observed in autodiploids: trisomy III (which fixes in five populations) and trisomy VIII (which fixes in one) (Table 2). CNVs are common in autodiploid genomes. Of the 46 autodiploid populations, CNVs appear in 19 and fix in 14. The 19 independently occurring autodiploid CNVs fall into 10 groups based on genomic position (Table 2). Autodiploid CNVs consist of both amplifications (n=4) and deletions (n=6). In contrast, no aneuploidies and only two amplifications are detected amongst the 40 haploid populations. These two amplifications are also observed in autodiploids.

**Table 2:** Structural variants in evolved autodiploids.

| Chr. | Start (kb) | End (kb) | Length (kb) | Copy Number | Description | Type | Clones* |
| --- | --- | --- | --- | --- | --- | --- | --- |
| I | 210 | 225 | 15 | 1N | CNV | loss | **B01a, B01b, E11a, E11b** |
| III | 85 | 85 | <10 kb | 0N | CNV | loss | **G01a, G01b, G01c** |
| III | 150 | 170 | 20 | 1N | CNV | loss | **A02a, A02b, B10a, B10b, C11a, C11b, C11c, F10a** |
| IV<sup>2</sup> | 900 | 1000 | 100 | 3N | CNV | gain | **B12a, B12b, C03a, E12a, E12b** |
| V<sup>3</sup> | 450 | 500 | 50 | 1N | CNV | loss | **B11a, B11b, F10a, F10b** |
| VIII | 525 | 545 | 20 | 1N | CNV | loss | **E11a, E11b** |
| XIII<sup>3</sup> | 190 | 200 | 10 | 1N | CNV | loss | C10a, D10a, E10c, H12a |
| XIII<sup>2</sup> | 190 | 200 | 10 | 3N | CNV | gain | **F02a, F02b** |
| XIV | 545 | 560 | 15 | 3N | CNV | gain | **A12a, A12b** |
| XV | 900 | 1100 | 200 | 3N | CNV | gain | G02b |
| III | 0 | 317 | 317 | 3N<sup>1</sup> | aneuploidy | gain | **C01a, C01b, D01a, D01b, D03a, D03b, E12a, E12b<sup>1</sup>, H02a, H02b,** |
| VIII | 0 | 924 | 924 | 3N | aneuploidy | gain | **A11a, A11b** |

\* Bolded clones indicate the CNV was found in all clones of the population
<sup>1</sup> Observed at 4N in one clone
<sup>2</sup> Also observed in one haploid
<sup>3</sup> Contains essential genes

### Autodiploids are buffered from deleterious mutations

To determine the extent to which an increase in ploidy buffers diploid lineages against the effects of deleterious mutations, we compared the frequency of mutations in essential genes in autodiploids with those of *MAT***a** haploids described previously (Lang *et al.* 2013). We specifically analyzed frameshift and nonsense mutations that would likely phenocopy the null mutants used to characterize genes as essential. Sixty-three of 66 high impact mutations in essential genes are heterozygous. For the remaining three mutations, zygosity is inconclusive due to low coverage (**Fig. S2B**). We find high impact mutations in essential genes to be exceptionally rare in haploids, with only a single case observed (Fig. 5A). In contrast, autodiploids contain a significantly higher proportion of high impact mutations in essential genes (*x*^*2*^ (1) = 20.32, *p <*0.0001). As expected, the proportion of low impact mutations within essential genes is consistent across ploidies (*x*^*2*^ (1) = 0.909, *p* = 0.339). Essential genes are also present within two of the large deletions observed in autodiploids (**Table 2**).

**Fig. 5.**
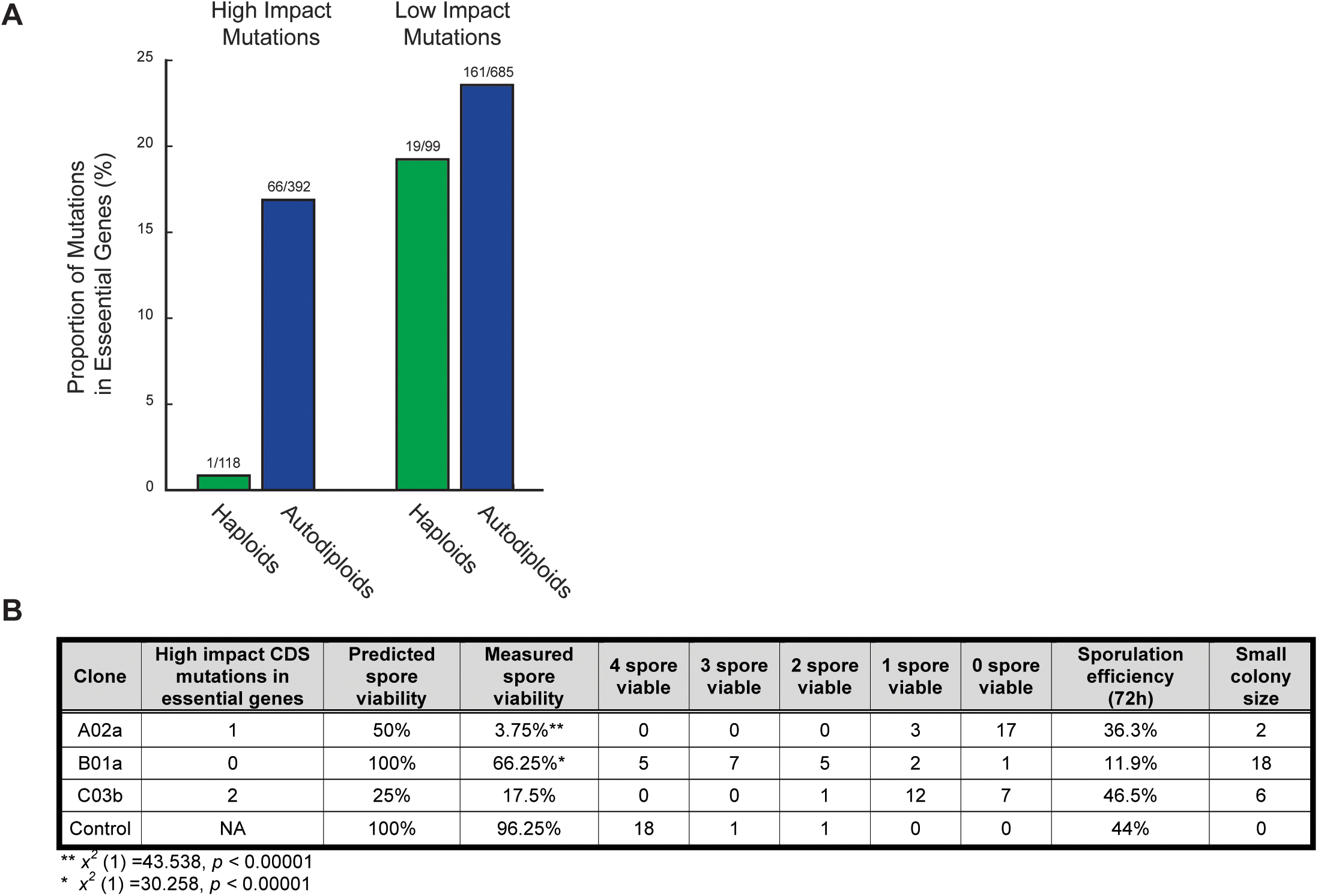
Recessive deleterious and lethal mutations. A) Shown are the proportions of high impact mutations (frameshift, nonsense) and low impact mutations (synonymous, intronic) in essential genes in haploids (green) and autodiploids (blue). Above each bar is the ratio of mutations in essential genes to mutations in all genes. B) Clones from three evolved diploid populations were sporulated and dissected. Spore viability and small colony size reflect recessive lethal and recessive deleterious mutations, respectively.

To experimentally validate that recessive lethal mutations accumulate in autodiploids, we sporulated three *MAT***a**/**a** from three different populations and performed tetrad dissections. Clones A02a, B01a, and C03b were selected because they contain no identifiable aneuploidies that would complicate measures of spore viability. Out of 20 total dissected tetrads (80 total spores) per clone, spore viability ranged from 4% to 66% in evolved autodiploid clones (Fig. 5B). Further, a substantial fraction of germinated spores developed morphologically small colony sizes relative to controls. We compared observed spore viability to expected viability based on the number of high impact mutations in genes annotated as essential. The only clone for which we observed four-spore viable tetrads, B01a, is also the only clone with no predicted recessive lethal mutations. Nonetheless, both A03a and B01a have significantly lower spore viability than expected (Fig. 5B). This in part may be due a genetic load imposed by segregating deleterious alleles. Consistent with our sequencing data, these data indicate that diploidy permits the accumulation of recessive lethal and deleterious mutations on a relatively short time scale.

## DISCUSSION

Whole genome duplications (WGDs) are significant evolutionary events that have profound impacts on genome evolution. Evidence of ancient whole-genome duplication events is found within lineages ancestral to most extant eukaryotic taxa (Jaillon *et al.* 2004; Meyer and Van de Peer 2005; Tang *et al.* 2008), including at least two WGDs in the vertebrate lineage (Dehal and Boore 2005), and a WGD approximately 100 mya in the *Saccharomyces* lineage (Kellis, Birren, Lander 2004; Wolfe and Shields 1997). In addition, the existence of numerous contemporary polyploid taxa suggests that genome duplication plays a role in short-term adaptive evolution (Van de Peer, Maere, Meyer 2009). Here, we show that experimental evolution of haploid *Saccharomyces cerevisiae* results in rapid and recurrent WGD. Clones with duplicated genomes arise early in all 46 populations and fix rapidly. We show that the fixation of autodiploids is due to a high rate of occurrence and a large fitness effect conferred by WGD.

Although the invasion and subsequent fixation of autodiploids in haploid-founded lineages has been reported before in yeast (see Table 1), a clear fitness advantage to diploidy has not always been evident. By employing a competitive growth assay, we demonstrate a relatively large fitness effect of a duplicated genome in our selective environment. A 3.6% fitness effect is substantial: in a recent study we quantified fitness effects of over 116 mutations from 11 evolved lineages in the same conditions, and only 9 conferred a fitness benefit greater than 3.6% (Buskirk, Peace, Lang 2017). The biological basis of this fitness advantage is unclear. However, there are several strong possibilities. Increased cell size, differential gene regulation, and a diploid-specific proteome (De Godoy *et al.* 2008; Galitski *et al.* 1999) may all contribute to the adaptive advantage of diploidy. More generally, environmental robustness is often associated with increases in ploidy (Van de Peer, Maere, Meyer 2009).

The recurrent and remarkably parallel manner in which autodiploids arise and fix points to not only a large fitness effect, but a high rate of occurrence. Our previous work has shown that parallel evolution is evident at the level of genetic pathway and even gene (Buskirk, Peace, Lang 2017; Marad and Lang 2017). However, the extent of the convergence observed here – where all 46 populations evolve to be autodiploids – is unprecedented in our experimental system. While it cannot be dismissed that some autodiploids were present in the founding inoculum, they are below our 1% detection limit. Autodiploids at this low of a frequency in the incoculum is not sufficient to explain the extent of fixation observed **(Fig. S5).** Simulations indicate the probability of an autodiploid lineage at 1% fixing in 46 out of 46 replicate populations is 2.5 × 10^−3^. Furthermore, given the common dynamics observed in populations of both mating types, autodiploids would have had to arisen in “jackpot” fashion and reach a similar frequency in the inocula of both mating-types. These data strongly support independent WGD events in replicate populations, suggesting a high background rate of duplication. This is consistent with the observation of frequent WGD in mutation accumulation lines (Lynch *et al.* 2008). Using a barcode-enrichment assay, Venkataram *et al.* (2016)) found that roughly half of all evolved clones with increased fitness that arose in a short-term enrichment experiment possessed no mutation apart from a WGD. The rate of WGD, therefore, is likely several-fold higher than the per base pair mutation rate.

Given the prevalence of autodiploids in the present evolution experiment, it is worth asking why autodiploids were not reported in a previous haploid evolution experiment in which ostensibly the identical strain and conditions were used (Lang *et al.* 2013). It is possible that in the prior experiment autodiploids did not fix or they could have fixed but were not detected. Despite conscious efforts to maintain identical selective environments, subtle differences in the conditions may exist given that evolution experiments were conducted years apart in different facilities. Indeed, inconsistency in the appearance of WGD across experiments and conditions is common in the field (Gorter *et al.* 2017; Voordeckers *et al.* 2015). Even subtle differences in the evolution conditions could shift the selective benefit of autodiploidy and yield population dynamics different from those seen here. Alternatively, it is possible that autodiploids did fix in the previous haploid evolution experiment but went undetected. The populations analyzed in the haploid study were part of a larger ~600 population experiment, and the 40 focal populations were selected based on the presence of a sterile phenotype. Mutations producing sterile phenotypes are predominantly adaptive and recessive loss-of-function (Lang, Murray, Botstein 2009). The presence of such beneficial mutations would have biased the selection of populations towards those retaining haploidy. We analyzed a subset of the remaining ~560 populations by DNA content staining and find that ~30% (3 of 10) of them appear autodiploid at generation 1,000, though this is still less than we report here. Further at least one of the forty sequenced populations (RMS1-E09, Lang *et al.*, 2013) which appeared to be an autodiploid based on the presence of a large number of mutations present at a frequency of 0.5, was confirmed as 2N through ploidy-staining.

The consequences of WGD are apparent on both the phenotypic and genotypic level. One such consequence is the susceptibility of autodiploids to Haldane’s sieve, resulting in a “depleted” spectrum of beneficial mutations. We find a decline in adaptation rate following WGD, which mirrors findings from studies that directly compare the rates of haploid population adaptation with that of diploids (Gerstein *et al.* 2011; Marad and Lang 2017). This implies a fitness tradeoff in the shift from 1N to 2N, wherein the fixation of a large-effect beneficial genotype comes at the cost of eliminating access to future recessive beneficial mutations. This tradeoff associated with genome duplication is predicted when population size is large and most beneficial mutations are partially or fully recessive (Otto 2007), conditions that are met in our populations (Lang *et al.* 2013; Marad and Lang 2017).

Autodiploids share physiological traits with both haploid and diploid cell types. Like their haploid founders, autodiploids possess only a single mating-type allele and will readily mate with cells of the opposite mating-type, indicating haploid-specific regulation of mating-pathway genes. As with diploids, autodiploids possess a 2N genome and exhibit larger cell size (Galitski *et al.* 1999). Consequently, we observe some overlap in the spectrum of beneficial mutations. We have identified targets of selection shared between haploids and autodiploids along with targets specific to autodiploids. While several targets were mutual to haploids and autodiploids, the extent of recurrence varied by gene. For example, *IRA1* mutations were common to both ploidies but enriched in haploids. In contrast, there were five ploidy-specific genes that were targets in autodiploids but never mutated in haploids. These genes (*PHO81, YTA7, PHO4, IRC8,* and *PSA1*) represent targets of selection that are specifically enriched in autodiploids, suggesting that WGD may expose adaptive pathways that are not easily accessible to either haploids or diploids.

Genome duplication also has consequences on genome stability and the evolution of structural variation. Across our 46 populations we identify 6 independently evolved aneuploidies and 20 independently evolved structural variants. Structural variants are more frequent in autodiploid genomes than in evolved haploid genomes of the same background, even after accounting for length of evolution. Haploids are constrained: whereas the structural variants observed in haploids always result in a net gain of genetic material, autodiploid structural variants include both amplifications and deletions. The ability to generate a greater degree of structural variation could provide a secondary advantage to WGD. Aneuploidies, large rearrangements, and CNVs have been shown to arise and confer an advantage in experimentally evolving yeast populations (Chang *et al.* 2013; Selmecki *et al.* 2015). Of note, several of the recurrent structural arrangements described in the present study, including trisomy III and a 317 kb deletion on Chr. III, are described as beneficial in Sunshine *et al.* (2015). The observation of both gain and loss of genetic material from Chr. III may indicate complex selection on phenotypes unachievable through point mutations.

Loss of heterozygosity (LOH) provides a means of overcoming the masking effect of ploidy in autodiploids allowing recessive beneficial mutations to become homozygous. Analysis of the distribution of homozygous mutations across evolved autodiploid genomes reveals LOH frequently occurs in two locations: on the right arm of Chr. XII and the right arm of Chr. XV. The right arm of Chr. XII has been characterized as a hotspot for LOH in experimental and natural populations (Magwene *et al.* 2011; Marad and Lang 2017) mediated by a high rate of recombination at the rDNA repeats (Keil and Roeder 1984). To our knowledge, a mitotic recombination hotspot on Chr. XV has not been described. Recurrent LOH may be have substantial evolutionary implications as the affected regions may experience different rates of genome evolution and divergence than the rest of the genome. On one hand, adaptation may be slow due to the periodic purging of variation and exposure of deleterious mutations to selection. On the other hand, the rate of adaptation may be increase by providing access to recessive beneficial mutations that would otherwise be masked by Haldane’s sieve. Theory predicts that sufficient mitotic recombination may allow asexual populations to circumvent Haldane’s sieve (Mandegar and Otto 2007). While we only show prevalence of LOH and not functional evidence of adaptive LOH, such events have been repeatedly observed in adapting yeast populations (Gerstein, Kuzmin, Otto 2014; Smukowski Heil *et al.* 2017). Further, the LOH on Chr. XV was not detected previously in diploids (Marad and Lang 2017), an observation that is more easily explained by selection than a change in the rate of occurrence.

The same masking effect that stifles recessive beneficial mutations is also predicted to permit the accumulation of deleterious mutations in diploids (Mable and Otto 2001). In evolved haploid populations few if any deleterious mutations fix: previously only 1 of 116 evolved mutations was characterized as putatively deleterious (Buskirk, Peace, Lang 2017). We show that, in contrast to haploid genomes, evolved autodiploid genomes harbor an abundance of putative recessive lethal mutations (Fig. 5A). We sporulated autodiploids with normal 2N karyotypes by complementing the *MAT*α information on a plasmid. We find evidence of the accumulation of both lethal and deleterious mutations as indicated by a large number of inviable and slow-growing haploid spores (Fig. 5B). The accumulation of recessive deleterious mutations in the genomes of clonal diploids may have long-term effects on adaptation. With each successive recessive deleterious mutation that fixes, genetic redundancy is eliminated, causing a shift in the distribution of fitness effects and an increase in the target size for lethal or deleterious mutations. Interestingly, loss of redundancy occurred rapidly following the historical yeast WGD (Scannell *et al.* 2006). Here we show that recessive deleterious and lethal mutations can accumulate shortly after WGD. On a population level, the increased target size for neutral mutations may increase standing variation between selective sweeps and may explain populations with deeply diverging clones (**Fig. S8**).

The ancient WGD in the *Saccharomyces* lineage is thought to have occurred by alloduplication followed by LOH at the mating-type locus to restore fertility (Marcet-Houben and Gabaldón 2015; Wolfe 2015), and therefore would have gone through an intermediate asexual ‘duplicated’ diploid state, similar to the *MAT***a**/**a** and *MAT*α/α populations investigated here. We demonstrate that this cell type has a direct fitness advantage over an isogenic haploid cell type. The immediate fitness gain of WGD is accompanied by several evolutionary tradeoffs that impact future adaptability including a reduced rate of adaptation, shifted distribution of beneficial mutations, karyotype changes, and the accumulation of recessive deleterious and lethal mutations that reduces redundancy in the duplicated genome.

## METHODS

### Strain construction

*MAT***a**/**a** strains were constructed for fitness assays by converting yGIL701, a fluorescently labeled *MAT***a**/α diploid isogenic to our ancestral haploid background, to *MAT***a**/**a**. yGIL701 was struck out and 10 separate clones were selected. Clones were transformed with pGIL088, which encodes a gal-inducible *HO* and a *MAT***a** specific *HIS3* marker. 5 ml cultures of YPD were inoculated with a single transformant for each starting clone. Cultures were grown for 48 hours, allowing for glucose to be depleted and catabolite repression of *GAL* genes to be lifted. After 48 hours 100 µl of each culture was plated to SD –his. Histidine prototrophs were screened in *α*-Factor (Sigma) for shmoos. Confirmed strains were used in competition assays.

### Evolution experiment

Experimental populations were founded with 130 µl of isogenic W303 ancestral culture; 22 with yGIL432 (*MAT***a**, *ade2-1*, *CAN1*, *his3-11*, *leu2-3,112*, *trp1-1*, *URA3*, *bar1*Δ::*ADE2*, *hm*lαΔ::*LEU2*, *GPA1*::NatMX, *ura3*Δ::P*FUS1*-yEVenus), and 24 with yGIL646, a *MAT*α strain otherwise isogenic to yGIL432. Populations analyzed here were evolved in separate wells of a 96-well plate. Ancestral strains were grown as 5 ml overnight cultures from single colonies prior to 96 well plate inoculation. This founding plate was propagated forward and then immediately frozen down.

All populations analyzed here were evolved in rich glucose (YPD) medium. Cultures were grown in unshaken 96-well plates at 30°C and were propagated every 24 hours via serial dilutions of 1:1024. Approximately every 60 generations, populations were cryogenically archived in 15% glycerol.

### Fitness assays

Fitness assays were performed as described previously (Buskirk, Peace, Lang 2017). Evolved autodiploid populations were mixed 1:1 with a version of the ancestral strain (yGIL432 or yGIL646, genotypes listed above) labeled with ymCitrine at *URA3.* Cultures were propagated in a 96-well plate in an identical fashion to the evolution experiment for 40 generations. Every 10 generations, saturated cultures were sampled for flow cytometry. Analysis of flow cytometry data was done using FlowJo 10.3. Selective coefficient was calculated as the slope of the change in the natural log ratio between query and reference strains. Assays were performed for all 46 evolved populations at 16 time points between generations 0 and 4,000.

To measure the fitness effect of autodiploidy, fitness assays were performed as described above, using instead a non-labeled version of yGIL432 as a reference. This strain was mixed 1:1 with either a fluorescently-labeled version of the same strain or one of ten biological replicate fluorescently labeled diploid strains. The fitness of each autodiploid reconstruction was calculated as the mean fitness across 12 replicate competitions.

Adaptation rates for each autodiploidized lineage were calculated as the rate of change between generation 0 and the time point at which diploids were present at over 98%. For comparison, rate of adaptation was also calculated for 39 diploid-founded populations evolved in parallel (Marad and Lang 2017). The median time point of autodiploid fixation was generation 600 for the haploid-founded dataset. To generate a comparable dataset, rates of adaptation for diploids were calculated from generations 0-600 and 600-4000. Rates were compared in SPSS using a repeated measures ANOVA with two within subject factors (time) and two between subject factors (haploid-founded and diploid-founded). Because some groups violated homogeneity assumptions, post-hoc analysis was done using a Bonferroni correction.

### DNA content analysis

Time-course ploidy states of 16 focal evolved populations were assayed through flow cytometry analysis of DNA content as described in Gerstein and Otto (2011). Briefly, 10 µl of each sample were inoculated in 3 ml YPD and grown overnight. 100 µl of saturated cultures were then diluted 1:50 into YPD and grown to mid-log. To arrest in G1, 1 ml mid-log culture was transferred into 200 µl 1M hydroxyurea and incubated on a 30°C roller drum for 3 hours. Cultures were then fixed with 70% ethanol, treated with RNAse and proteinase K, stained with Cytox green (Molecular Probes), and analyzed on a BD FACSCanto. Haploid and diploid frequencies were estimated using FlowJo v10.3 by fitting data to Watson-Pragmatic cell cycle models. This method of estimation was validated with a series of known ploidy mixtures (**Fig. S9**).

### Simulations

Simulations of lineage trajectories were performed using a forward-time algorithm designed to imitate the same conditions as our evolution experiment. Estimates for the distribution of fitness effects (an exponential distribution with mean 
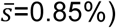
 and beneficial mutation rate (*U*_*b*_ =1.0 x 10^−4^) were described previously for our experimental conditions (Frenkel, Good, Desai 2014). This model assumes the spectrum of mutations available to haploids is the same as the spectrum available to autodiploids. Simulations were performed with constant inputs for DFE parameters, beneficial mutation rate, inoculation time of the focal lineage (generation *t* =0), and fitness advantage of the focal lineage (*s*_*0*_ =3.6%). The initial frequency of the focal lineage was varied (*f*_*0*_ = 0.01%-1.0%) for each set of simulations, and a total of ten thousand simulations were performed for each *f*_*0*_.

### Sequencing

Evolved clones were obtained by streaking evolved populations to singles on YPD and selecting two clones per population. These clones were grown to saturation in 5 ml YPD and then spun down to cell pellets and frozen at −20°C. Genomic DNA was harvested from frozen pellets via phenol-chloroform extraction and precipitated in ethanol. Total genomic DNA was used in a Nextera library preparation. The Nextera protocol was followed as described previously (Buskirk, Peace, Lang 2017). All individually barcoded clones were pooled and sequenced on 2 lanes of an Illumina HiSeq 2500 sequencer by the Sequencing Core Facility at the Lewis-Sigler Institute for Integrative Genomics at Princeton.

### Sequencing analysis

Two lanes of raw sequence data were concatenated and then demultiplexed using a custom python script (barcodesplitter.py) from L. Parsons (Princeton University). Adapter sequences were trimmed using the fastx_clipper from the FASTX Toolkit. Trimmed reads were aligned to an S288c reference genome version R64-2-1 (Engel and Cherry 2013) using BWA v0.7.12 (Li and Durbin 2009) and variants were called using FreeBayes v0.9.21-24-381 g840b412 (Garrison and Marth 2012). Roughly 10,000 polymorphisms were detected between our ancestral W303 background and the S288c reference, and the corresponding genomic positions were removed from analysis. All remaining calls were confirmed manually by viewing BAM files in IGV (Thorvaldsdóttir, Robinson, Mesirov 2013). Zygosity was determined based on read depth and allele frequency (**Fig. S2B**). Mutations were classified as fixed if present in all clones from a population. Clones were genotyped for *MAT* alleles by identifying mating-type specific sequences within the demultiplexed FASTQ files.

Clone genomes were each independently queried for structural variants. Following BWA alignment, coverage at each position across the genome was calculated. Aneuploidies were detected by calculating median chromosome coverage and dividing this by median genome-wide coverage for each chromosome, producing an approximate chromosome copy number relative to the duplicated genome (**Fig. 4; Dataset 2**). CNVs were detected by visual inspection of chromosome coverage plots created in R (**Fig. S10; Dataset 3**).

**Fig. 4.**
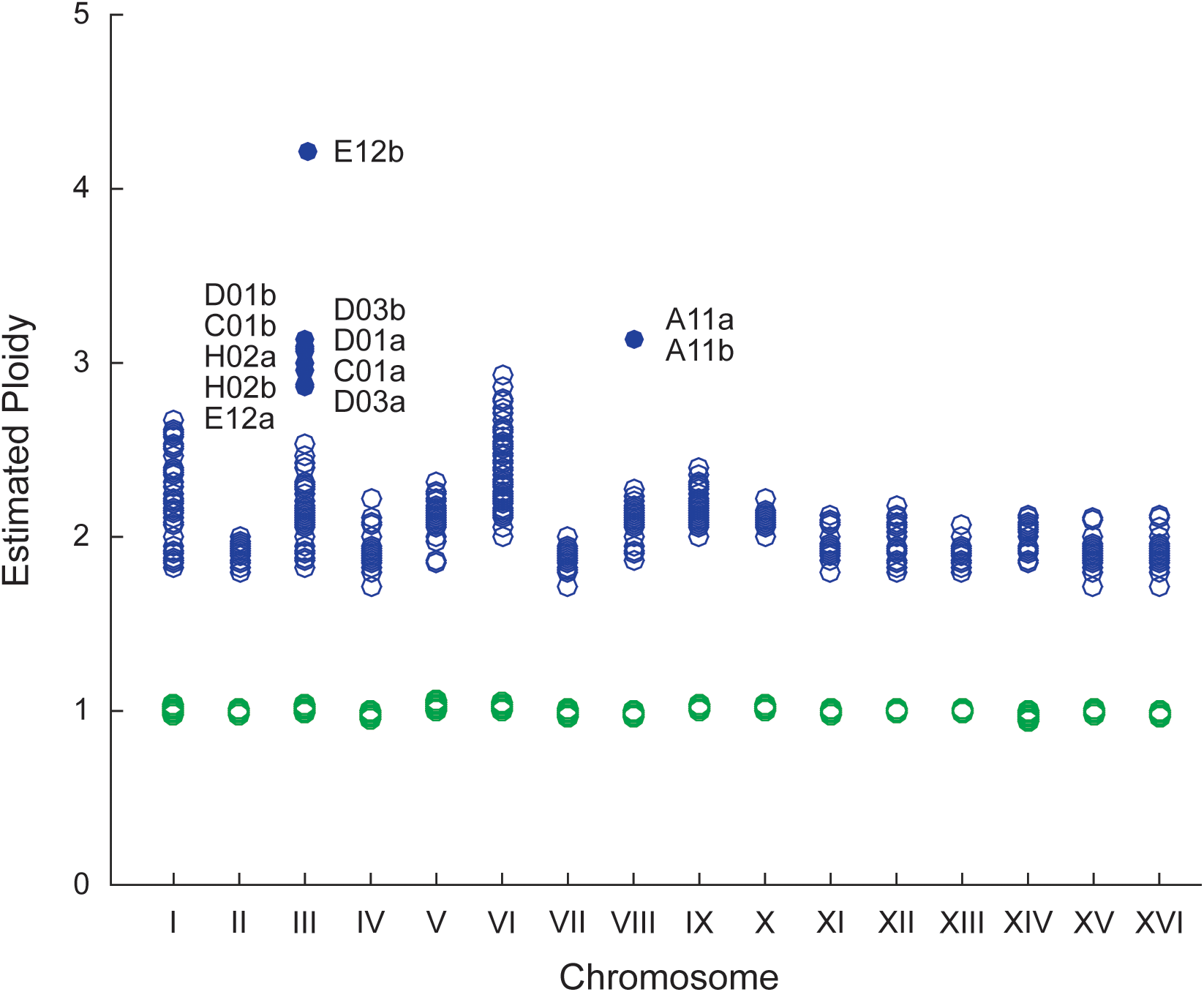
Detection of Aneuploidies. For each sequenced sample, coverage across each chromosome was compared to genome-wide coverage. Based on DNA content staining, baseline ploidy was assumed to be 1N for haploids and 2N for autodiploids. Euploidy is indicated by empty circles: haploid - green, autodiploids - blue. Aneuploidies are shown as filled circles and labeled by clone.

### Phylogenetic analysis

Variants identified by SNPeff were used to infer a phylogeny based on 7,932 sites containing 4,742 variable sites, either SNPs or small indels (**Fig. S8**). Evolved and ancestral sequences (n=93) were aligned with MUSCLE. A general time reversible substitution model with uniform rates (-lnL= 44803.45) was selected based on jModelTest. A maximum likelihood tree was then constructed and rooted by the ancestor in MEGA. Subclades were found to be due to incomplete linage sorting of mitochondrial polymorphisms. After phylogenetic analysis it was evident that four clones were originally attributed to incorrect populations. Tight clustering and short branch lengths suggests either very recent contamination or an issue during colony isolation (populations were struck out two to a plate on bisected YPD plates). In the text, these clones are identified by the suffix “c” and are attributed to the population to which they are most phylogenetically similar.

### Identification of common targets and ploidy-enriched targets

A recurrence approach was utilized to identify common targets of selection. A random distribution of the 3,431 coding sequence (CDS) mutations across all 5,800 genes predicts only two genes to be mutated more than five times by chance alone. We determined the probability that chance alone explains the observed number of mutations of each gene by assuming a random distribution of the 3,431 mutations across the 8,527,393 bp genome-wide CDS. Common targets of selection were defined as genes with five or more CDS mutations and a corresponding probability of less than 0.1% (Fig. 2A). Notably, analysis using only nonsynonymous mutations identified largely the same set of common targets of selection as did analysis using all CDS mutations. To determine which targets of selection are impacted by ploidy, our recurrence approach was used to analyze mutations in a previously published *MAT***a** haploid dataset (**Fig. S7**) (Lang *et al.* 2013). We compared the probability of the observed number of CDS mutations in each gene between ploidies (Fig. 2C). A gene was considered ploidy-enriched if the ratio of probabilities was at least 10^5^.

### Evolved clone sporulation and tetrad dissection

Three clones (A02a, B01a, C03b) for which genome sequence data revealed no aneuploidies were selected for sporulation. Evolved *MAT***a**/**a** clones were transformed with pGIL071 which encodes the *α*2 gene necessary for sporulation and a *URA3* marker for selection. Transformants were sporulated in Spo++ -ura media. Following 72 hours, sporulation efficiency was calculated via hemocytometer, cultures were digested with zymolyase, and tetrads were dissected on YPD agar plates. Spores were incubated 48 hours and then assayed for germination. Control strain yGIL1039, made by crossing yGIL432 to yGIL646 and converting the resulting diploid to *MAT***a/a** as described above, was transformed and dissected in parallel.

## AUTHOR CONTRIBUTIONS

KJF, SWB, and GIL conceived of the project and designed experiments. KJF and SWB performed library preparations and SWB performed sequencing analysis and bioinformatics. KJF performed the experiments. KJF, DAM, and RCV collected time-course ploidy data. RCV performed simulations. DAM collected time-course fitness data. KJF, SWB, and GIL analyzed data and wrote the manuscript.

## ACKNOWLEDGEMENTS

We thank Alex Nguyen (Desai lab, Harvard) for providing the plasmid with mating-type specific markers. We thank Aleeza Gerstein and Jun-Yi Leu for comments on the manuscript. This work was supported by the Charles E. Kaufman Foundation of The Pittsburgh Foundation.

## DATA DEPOSITION

The short-read sequencing data reported in this paper have been deposited in the NCBI BioProject database (accession no. PRJNA422100).

## SUPPLEMENTAL DATASETS

**Dataset 1:** All 8,305 de novo mutations detected across the 46 autodiploid populations

**Dataset 2:** Aneuploidies detected by sequencing read depth

**Dataset 3:** Copy number variants detected by sequencing read depts

## REFERENCES

Buskirk SW, Peace RE, Lang GI. 2017. Hitchhiking and epistasis give rise to cohort dynamics in adapting populations. Proc Natl Acad Sci U S A 114(31):8330–5.

Chang S, Lai H, Tung S, Leu J.2013. Dynamic large-scale chromosomal rearrangements fuel rapid adaptation in yeast populations. PLoS Genetics 9(1):e1003232.

Cooper VS, Schneider D, Blot M, Lenski RE. 2001. Mechanisms causing rapid and parallel losses of ribose catabolism in evolving populations of *Escherichia coli B*. J Bacteriol 183(9):2834–41.

De Godoy LM, Olsen JV, Cox J, Nielsen ML, Hubner NC, Fröhlich F, Walther TC, Mann M.2008. Comprehensive mass-spectrometry-based proteome quantification of haploid versus diploid yeast. Nature 455(7217):1251–4.

Dehal Pand Boore JL. 2005. Two rounds of whole genome duplication in the ancestral vertebrate. PLoS Biology 3(10):e314.

Deutschbauer AM, Jaramillo DF, Proctor M, Kumm J, Hillenmeyer ME, Davis RW, Nislow C, Giaever G.2005. Mechanisms of haploinsufficiency revealed by genome-wide profiling in yeast. Genetics 169(4):1915–25.

Dunham MJ, Badrane H, Ferea T, Adams J, Brown PO, Rosenzweig F, Botstein D.2002. Characteristic genome rearrangements in experimental evolution of *Saccharomyces cerevisiae*. Proc Natl Acad Sci U S A 99(25):16144–9.

Engel SRand Cherry JM. 2013. The new modern era of yeast genomics: Community sequencing and the resulting annotation of multiple *Saccharomyces cerevisiae strains at the Saccharomyces genome database*. Database 2013:bat012.

Ezov TK, Boger-Nadjar E, Frenkel Z, Katsperovski I, Kemeny S, Nevo E, Korol A, Kashi Y.2006. Molecular‐ genetic biodiversity in a natural population of the yeast *Saccharomyces cerevisiae* from “Evolution Canyon”: Microsatellite polymorphism, ploidy and controversial sexual status. Genetics 174(3):1455–68.

Frenkel EM, Good BH, Desai MM. 2014. The fates of mutant lineages and the distribution of fitness effects of beneficial mutations in laboratory budding yeast populations. Genetics 196(4):1217–26.

Galitski T, Saldanha AJ, Styles CA, Lander ES, Fink GR. 1999. Ploidy regulation of gene expression. Science 285(5425):251–4.

Garrison Eand Marth G.2012. Haplotype-based variant detection from short-read sequencing. arXiv Preprint arXiv:1207.3907

Gerstein A, Kuzmin A, Otto S.2014. Loss-of-heterozygosity facilitates passage through Haldane's sieve for *Saccharomyces cerevisiae* undergoing adaptation. Nature Communications 5:3819.

Gerstein ACand Otto SP. 2011. Cryptic fitness advantage: Diploids invade haploid populations despite lacking any apparent advantage as measured by standard fitness assays. PLoS One 6(12):e26599.

Gerstein AC, Lim H, Berman J, Hickman MA. 2017. Ploidy tug‐of‐war: Evolutionary and genetic environments influence the rate of ploidy drive in a human fungal pathogen. Evolution 71(4):1025–38.

Gerstein AC, Cleathero L, Mandegar M, Otto S.2011. Haploids adapt faster than diploids across a range of environments. J Evol Biol 24(3):531–40.

Gerstein AC, Chun HE, Grant A, Otto SP. 2006. Genomic convergence toward diploidy in *Saccharomyces cerevisiae*. PLoS Genetics 2(9):e145.

Gerstein AC. 2013. Mutational effects depend on ploidy level: All else is not equal. Biol. Lett. 9(1):20120614.

Gorter FA, Derks MFL, van den Heuvel J, Aarts MGM, Zwaan BJ, de Ridder D, de Visser JAGM. 2017. Genomics of adaptation depends on the rate of environmental change in experimental yeast populations. Molecular Biology and Evolution 34(10):2613–26.

Gregory TR. 2001. Coincidence, coevolution, or causation? DNA content, cell size, and the C-value enigma. Biological Reviews 76(1):65–101.

Gresham D, Desai MM, Tucker CM, Jenq HT, Pai DA, Ward A, DeSevo CG, Botstein D, Dunham MJ. 2008. The repertoire and dynamics of evolutionary adaptations to controlled nutrient-limited environments in yeast. PLoS Genet 4(12):e1000303.

Hong Jand Gresham D.2014. Molecular specificity, convergence and constraint shape adaptive evolution in nutrient-poor environments. PLoS Genetics 10(1):e1004041.

Jaillon O, Aury J, Brunet F, Petit J, Stange-Thomann N, Mauceli E, Bouneau L, Fischer C, Ozouf-Costaz C, Bernot A.2004. Genome duplication in the teleost fish *Tetraodon nigroviridis* reveals the early vertebrate proto-karyotype. Nature 431(7011):946–57.

Keil RLand Roeder GS. 1984. *Cis*-acting, recombination-stimulating activity in a fragment of the ribosomal DNA of *S. cerevisiae*. Cell 39(2):377–86.

Kellis M, Birren BW, Lander ES. 2004. Proof and evolutionary analysis of ancient genome duplication in the yeast *Saccharomyces cerevisiae*. Nature 428(6983):617–24.

Kosheleva Kand Desai MM. 2017. Recombination alters the dynamics of adaptation on standing variation in laboratory yeast populations. Mol Biol Evol. Jan 1;35(1):180–201.

Kvitek DJand Sherlock G.2013. Whole genome, whole population sequencing reveals that loss of signaling networks is the major adaptive strategy in a constant environment. PLoS Genetics 9(11):e1003972.

Lang GI, Murray AW, Botstein D.2009. The cost of gene expression underlies a fitness trade-off in yeast. Proc Natl Acad Sci U S A 106(14):5755–60.

Lang GI, Rice DP, Hickman MJ, Sodergren E, Weinstock GM, Botstein D, Desai MM. 2013. Pervasive genetic hitchhiking and clonal interference in forty evolving yeast populations. Nature 500(7464):571–4.

Lenski REand Travisano M.1994. Dynamics of adaptation and diversification: A 10,000-generation experiment with bacterial populations. Proc Natl Acad Sci U S A 91(15):6808–14.

Li Hand Durbin R.2009. Fast and accurate short read alignment with Burrows–Wheeler transform. Bioinformatics 25(14):1754–60.

Liti G.2015. The natural history of model organisms: The fascinating and secret wild life of the budding yeast *S. cerevisiae*. Elife 4:e05835.

Lynch M, Sung W, Morris K, Coffey N, Landry CR, Dopman EB, Dickinson WJ, Okamoto K, Kulkarni S,Hartl DL, et al.2008. A genome-wide view of the spectrum of spontaneous mutations in yeast. Proc Natl Acad Sci U S A 105(27):9272–7.

Mable BKand Otto SP. 2001. Masking and purging mutations following EMS treatment in haploid, diploid and tetraploid yeast (*Saccharomyces cerevisiae*). Genetics Research 77(1):9–26.

Magwene PM, Kayikci O, Granek JA, Reininga JM, Scholl Z, Murray D.2011. Outcrossing, mitotic recombination, and life‐history trade-offs shape genome evolution in *Saccharomyces cerevisiae*. Proc Natl Acad Sci U S A 108(5):1987–92.

Mandegar MAand Otto SP. 2007. Mitotic recombination counteracts the benefits of genetic segregation. Proc Biol Sci 274(1615):1301–7.

Marad DAand Lang GI. 2017. Restricted access to beneficial mutations slows adaptation and biases fixed mutations in diploids. bioRxiv: 171462

Marcet-Houben Mand Gabaldón T.2015. Beyond the whole-genome duplication: Phylogenetic evidence for an ancient interspecies hybridization in the baker’s yeast lineage. PLoS Biology 13(8):e1002220.

Meyer Aand Van de Peer Y.2005. From 2R to 3R: Evidence for a fish‐specific genome duplication (FSGD). Bioessays 27(9):937–45.

Orr HAand Otto SP. 1994. Does diploidy increase the rate of adaptation?Genetics 136(4):1475–80.

Otto SP. 2007. The evolutionary consequences of polyploidy. Cell 131(3):452–62.

Oud B, Guadalupe-Medina V, Nijkamp JF, de Ridder D, Pronk JT, van Maris AJ, Daran JM. 2013. Genome duplication and mutations in *ACE2* cause multicellular, fast-sedimenting phenotypes in evolved *Saccharomyces cerevisiae*. Proc Natl Acad Sci U S A 110(45):E4223–31.

Scannell DR, Byrne KP, Gordon JL, Wong S, Wolfe KH. 2006. Multiple rounds of speciation associated with reciprocal gene loss in polyploid yeasts. Nature 440(7082):341–5.

Sellis D, Callahan BJ, Petrov DA, Messer PW. 2011. Heterozygote advantage as a natural consequence of adaptation in diploids. Proc Natl Acad Sci U S A 108(51):20666–71.

Selmecki AM, Maruvka YE, Richmond PA, Guillet M, Shoresh N, Sorenson AL, De S, Kishony R, Michor F, Dowell R.2015. Polyploidy can drive rapid adaptation in yeast. Nature 519(7543):349–52.

Smukowski Heil CS, DeSevo CG, Pai DA, Tucker CM, Hoang ML, Dunham MJ. 2017. Loss of heterozygosity drives adaptation in hybrid yeast. Mol Biol Evol 34(7):1596–612.

Sunshine AB, Payen C, Ong GT, Liachko I, Tan KM, Dunham MJ. 2015. The fitness consequences of aneuploidy are driven by condition-dependent gene effects. PLoS Biology 13(5):e1002155.

Tang H, Bowers JE, Wang X, Ming R, Alam M, Paterson AH. 2008. Synteny and collinearity in plant genomes. Science 320(5875):486–8.

Thorvaldsdóttir H, Robinson JT, Mesirov JP. 2013. Integrative genomics viewer (IGV): High-performance genomics data visualization and exploration. Briefings in Bioinformatics 14(2):178–92.

Van de Peer Y, Maere S, Meyer A.2009. The evolutionary significance of ancient genome duplications. Nature Reviews Genetics 10(10):725–32.

Venkataram S, Dunn B, Li Y, Agarwala A, Chang J, Ebel ER, Geiler-Samerotte K, Hérissant L, Blundell JR, Levy SF. 2016. Development of a comprehensive genotype-to-fitness map of adaptation-driving mutations in yeast. Cell 166(6):1585,1596. e22.

Voordeckers K, Kominek J, Das A, Espinosa-Cantú A, De Maeyer D, Arslan A, Van Pee M, van der Zande E, Meert W, Yang Y.2015. Adaptation to high ethanol reveals complex evolutionary pathways. PLoS Genetics 11(11):e1005635.

Wolfe KH. 2015. Origin of the yeast whole-genome duplication. PLoS Biology 13(8):e1002221.

Wolfe KHand Shields DC. 1997. Molecular evidence for an ancient duplication of the entire yeast genome. Nature 387(6634):708.

Zeyl C, Vanderford T, Carter M.2003. An evolutionary advantage of haploidy in large yeast populations. Science 299(5606):555–8.

Zhang H, Zeidler AF, Song W, Puccia CM, Malc E, Greenwell PW, Mieczkowski PA, Petes TD, Argueso JL. 2013. Gene copy-number variation in haploid and diploid strains of the yeast *Saccharomyces cerevisiae*. Genetics 193(3):785–801.

Zörgö E, Chwialkowska K, Gjuvsland AB, Garré E, Sunnerhagen P, Liti G, Blomberg A, Omholt SW, Warringer J.2013. Ancient evolutionary trade-offs between yeast ploidy states. PLoS Genetics 9(3):e1003388.

